# Modeling semantic encoding in a common neural representational space

**DOI:** 10.1101/288605

**Authors:** Cara E. Van Uden, Samuel A. Nastase, Andrew C. Connolly, Ma Feilong, Isabella Hansen, M. Ida Gobbini, James V. Haxby

## Abstract

Encoding models for mapping voxelwise semantic tuning are typically estimated separately for each individual, limiting their generalizability. In the current report, we develop a method for estimating semantic encoding models that generalize across individuals. Functional MRI was used to measure brain responses while participants freely viewed a naturalistic audiovisual movie. Word embeddings capturing agent-, action-, object-, and scene-related semantic content were assigned to each imaging volume based on an annotation of the film. We constructed both conventional within-subject semantic encoding models and between-subject models where the model was trained on a subset of participants and validated on a left-out participant. Between-subject models were trained using cortical surface-based anatomical normalization or surface-based whole-cortex hyperalignment. We used hyperalignment to project group data into an individual’s unique anatomical space via a common representational space, thus leveraging a larger volume of data for out-of-sample prediction while preserving the individual’s fine-grained functional–anatomical idiosyncrasies. Our findings demonstrate that anatomical normalization degrades the spatial specificity of between-subject encoding models relative to within-subject models. Hyperalignment, on the other hand, recovers the spatial specificity of semantic tuning lost during anatomical normalization, and yields model performance exceeding that of within-subject models.

## Introduction

Recent neuroimaging work has revealed widespread cortical representation of semantic content conveyed by visual and linguistic stimuli (Huth et al., 2012, 2016; Pereira et al., 2018; Wehbe et al., 2014). These findings hinge on the development of forward encoding models, which find a mapping from stimuli to voxelwise responses via a complex intermediate feature space (Naselaris et al., 2011). These feature spaces may capture distributional properties of large corpora of text (e.g., word co-occurrence) in the case of semantic representation (e.g., Huth et al., 2016; Mitchell et al., 2008), or comprise neurally-inspired models of vision (e.g., Güçlü and van Gerven, 2015; Kay et al., 2008; Nishimoto et al., 2011) or audition (e.g., de Heer et al., 2017; Santoro et al., 2014). If the intermediate feature space adequately captures stimulus qualities of interest and the model is trained on a sufficiently diverse sample of stimuli, the estimated model will generalize well to novel stimuli. Naturalistic stimuli and tasks (such as watching movies, listening to stories) enhance this approach by evoking reliable neural responses (Hasson et al., 2010) and broadly sampling stimulus space (Haxby et al., 2014), as well as increasing ecological validity (Felsen and Dan, 2005) and participant engagement (Vanderwal et al., 2017).

Although encoding models provide a fine-grained voxel-specific measure of functional tuning, they are typically estimated independently for each participant (e.g., Huth et al., 2012, 2016). This is problematic because we can collect only a limited volume of data in any one participant, and each participant’s model has limited generalizability across individuals (cf. Güçlü and van Gerven, 2017; Vodrahalli et al., 2017; Yamada et al., 2015). Recent work has demonstrated that group-level estimates of functional organization obscure marked individual-specific idiosyncrasies (Braga and Buckner, 2017; Gordon et al., 2017; Laumann et al., 2015). This is because functional–anatomical correspondence—the mapping between functional tuning and macroanatomical structure—varies considerably across individuals (Aine et al., 1996; Frost and Goebel, 2012; Riddle and Purves, 1995; Watson et al., 1993; Zhen et al., 2015, 2017). While macroanatomical normalization (i.e., nonlinear volumetric or cortical surface-based alignment) may be sufficient for capturing commonalities in coarse-grained functional areas, it cannot in principle align fine-grained functional topographies across individuals (cf. Conroy et al., 2013; Sabuncu et al., 2010). If we hope to predict functional tuning across individuals at the specificity of individual cortical vertices, we need to circumvent the correspondence problem between function and anatomy (Dubois and Adolphs, 2016; Poldrack, 2017).

In the following, we outline an approach for estimating encoding models that can make detailed predictions of responses to novel stimuli in novel individuals at the specificity of cortical vertices. To accommodate idiosyncratic functional topographies, we use hyperalignment to derive transformations to map each individual’s responses into a common representational space (Guntupalli et al., 2016; Haxby et al., 2011). The searchlight hyperalignment algorithm learns a locally-constrained whole-cortex transformation rotating each individual’s anatomical coordinate space into a common space that optimizes the correspondence of representational geometry (in this case, the response patterns to the movie stimulus at each time point) across brains. We use a dynamic, naturalistic stimulus—the *Life* nature documentary narrated by David Attenborough—for the dual purpose of deriving hyperalignment transformations and fitting the encoding model. Using a naturalistic paradigm that thoroughly samples both stimulus space and neural response space is critical for robustly fitting the encoding model and ensuring the hyperalignment transformations generalize to novel experimental contexts (Guntupalli et al., 2016; Haxby et al., 2011).

Although hyperalignment dramatically improves between-subject decoding (Guntupalli et al., 2016; Haxby et al., 2011), relatively few attempts have been made to integrate hyperalignment and voxelwise encoding models. Yamada and colleagues (2015) used a many-to-one sparse regression to predict voxel responses to simple visual stimuli across pairs of participants. Bilenko and Gallant (2016) implemented hyperalignment using regularized kernel canonical correlation analysis (Xu et al., 2012) to compare encoding models across subjects. Recent work (Vodrahalli et al., 2017) using a probabilistic, reduced-dimension variant of hyperalignment (Chen et al., 2015) has suggested that encoding models perform better in a lower-dimensional shared response space. Finally, Güçlü and van Gerven (2017) have employed hyperalignment in conjunction with a deep convolutional neural network (Tran et al., 2015) to predict responses to video clips in six dorsal stream visual areas (Nishimoto et al., 2011). They demonstrated that estimating an encoding model in a common representational space does not diminish model performance, and that aggregating additional subjects in the common spaces can improve performance.

To evaluate hyperalignment in the context of encoding models, we compared within-subject encoding models and between-subject encoding models where a model trained on three-fourths of the movie in a subset of participants is used to predict responses at each cortical vertex for the left-out fourth of the movie in a left-out participant. We compared between-subject models using high-performing surface-based anatomical normalization (Fischl, 2012; Klein et al., 2010) and surface-based searchlight whole-cortex hyperalignment (Guntupalli et al., 2016). We model semantic tuning at each cortical vertex based on distributed word embeddings (word2vec; Mikolov et al., 2013) assigned to each imaging volume based on an annotation of the documentary. We first show that constructing between-subject models using anatomical alignment reduces the spatial specificity of vertex-wise semantic tuning relative to within-subject models. Next, we demonstrate that hyperalignment generally leads to improved between-subject model performance, exceeding within-subject models. Hyperalignment effectively recovers the specificity of within-subject models, allowing us to leverage a large volume of group data for individualized prediction at the specificity of individual voxels or cortical vertices.

## Materials and methods

### Participants

Eighteen right-handed adults (10 female) with normal or corrected-to-normal vision participated in the experiment. Participants reported no neurological conditions. All participants gave written, informed consent prior to participating in the study, and the study was approved by the Institutional Review Board of Dartmouth College. These data have been previously used for the purpose of hyperalignment in a published report by Nastase and colleagues (2017).

### Stimuli and design

Participants freely viewed four segments of the *Life* nature documentary narrated by David Attenborough. The four runs were of similar duration (15.3 min, 14 min, 15.4 min, and 16.5 min), totaling 63 minutes. The movie stimulus included both the visual and auditory tracks, and sound was adjusted to a comfortable level for each participant. The video was back-projected on a screen placed at the rear of the scanner bore, and was viewed with a mirror attached to the head coil. Audio was delivered using MRI-compatible fiber-optic electrodynamic headphones (MR confon GmbH, Magdeburg, Germany). Participants were instructed to remain still and watch the documentary as though they were watching a movie at home. Note that this free viewing task contrasts with prior forward-encoding studies that enforced central fixation while viewing videos (e.g., Huth et al. 2012; Nishimoto et al. 2011), which we expect to affect the comparative performance of forward encoding models, especially in early visual cortex; however, a full treatment of the magnitude of such effects is beyond the scope of this paper. Stimuli were presented using PsychoPy (Peirce, 2007).

### Image acquisition

Structural and functional images were acquired using a 3T Philips Intera Achieva MRI scanner (Philips Medical Systems, Bothell, WA) with a 32-channel phased-array SENSE (SENSitivity Encoding) head coil. Functional, blood-oxygenation-level-dependent (BOLD) images were acquired in an interleaved fashion using single-shot gradient-echo echo-planar imaging with fat suppression and a SENSE parallel acceleration factor of 2: TR/TE = 2500/35 ms, flip angle = 90°, resolution = 3 mm^3^ isotropic voxels, matrix size = 80 × 80, FOV = 240 × 240 mm^2^, 42 transverse slices with full brain coverage and no gap, anterior–posterior phase encoding. Four runs were collected for each participant, consisting of 374, 346, 377, and 412 dynamic scans, or 935 s, 865 s, 942.5 s, and 1030 s, respectively. A T1-weighted structural scan was obtained using a high-resolution 3D turbo field echo sequence: TR/TE = 8.2/3.7 ms, flip angle = 8°, resolution = 0.9375 × 0.9375 × 1.0 mm^3^, matrix size = 256 × 256 × 220, and FOV = 240 × 240 × 220 mm^3^.

### Preprocessing

Raw data were organized to conform to the Brain Imaging Data Structure (BIDS; Gorgolewski et al., 2016) specifications and were preprocessed using fmriprep (Esteban et al., 2017; Gorgolewski et al., 2011, 2017), which provides a streamlined, state-of-the-art preprocessing pipeline that incorporates various software packages. Within the fmriprep framework, cortical surfaces were reconstructed from the T1-weighted structural images using FreeSurfer (Dale et al., 1999) and spatially normalized to the fsaverage6 template based on sulcal curvature (Fischl et al., 1999). Prior to spatial normalization, T2*-weighted functional volumes were slice-time corrected (Cox, 1996), realigned for head motion (Jenkinson et al., 2002), aligned to the anatomical image (Greve and Fischl, 2009), and sampled to the cortical surface. Time-series data were detrended using AFNI’s 3dTproject (Cox, 1996), which removes nuisance variables and trends via a single linear regression model. The regression model included a framewise displacement regressor (Power et al., 2012), the first six principal components from cerebrospinal fluid (Behzadi et al., 2007), head motion parameters, first- and second-order polynomial trends, and a band-pass filter (0.00667 to 0.1 Hz). All surface data were visualized in SUMA (Saad et al., 2004).

### Whole-brain hyperalignment

Surface-based searchlight whole-cortex hyperalignment (Guntupalli et al., 2016; Haxby et al., 2011) was performed based on the data collected while participants viewed the *Life* nature documentary using leave-one-run-out cross-validation: three of four runs were used to estimate the hyperalignment transformations for all participants; these transformations were then applied to the left-out run for model evaluation. The hyperalignment algorithm, described in detail by Guntupalli and colleagues (Guntupalli et al., 2016, 2018), uses iterative pairwise applications of the Procrustes transformation (Gower, 1975), effectively rotating a given subject’s multivariate response space to best align their patterns of response to time-points in the movie with a reference time series of response patterns.

In the first iteration, the response trajectory of an arbitrarily-chosen subject serves as the reference, and a second subject’s data are rotated via the Procrustes transformation into alignment with that reference. For each additional subject, a new reference trajectory is computed by averaging the previously aligned subject’s data and the reference, and the new subject is aligned to this reference. Aligning and averaging all subjects’ data in this way results in an intermediate template. In the second iteration, each subject’s data is again aligned to this intermediate reference, and the average of all subjects’ aligned response vectors are recomputed. This average response trajectory serves as the final functional template in a common representational space. For each subject, we calculate a final transformation to this functional template. These hyperalignment transformations can then be used to project data from a left-out run into the common representational space, or the transpose of a given subject’s transformation matrix can be used to project from the common space into a particular subject’s response space.

To locally constrain hyperalignment, we compute these transformations separately within large 20 mm radius surface-based searchlight disks centered on each cortical vertex (Guntupalli et al., 2016; Kriegeskorte et al., 2006; Oosterhof et al., 2011). Each searchlight comprised on average 610 vertices (*SD* = 162, median = 594, range: 237–1,238 vertices). The resulting rotation parameters are only defined for vertices within a given searchlight; however, searchlights are heavily overlapping. These local transformations are aggregated by summing overlapping searchlight transformation parameters to construct a single sparse transformation for each cortical hemisphere. For each subject, this results in two *N* × *N* transformation matrices, one for each cortical hemisphere, where *N* is the number of vertices in a hemisphere (40,962 for the fsaverage6 template). These matrices contain non-zero values only for vertices within the radius of a searchlight. Because each vertex is a constituent of many overlapping searchlights, the final rotation parameters for a given vertex will reflect transformations for all searchlights to which it contributes. Response time series for each vertex are z-scored before and after each application of the Procrustes transformation. All functional data were anatomically normalized to the fsaverage6 template prior to hyperalignment. This procedure is not strictly necessary for hyperalignment (and is not optimal due to interpolation during surface projection), but is used here for simplicity and to facilitate comparison between anatomically normalized and hyperaligned data. Note that hyperalignment does not yield a one-to-one mapping between voxels or vertices across subjects, but rather models each voxel’s or vertex’s response profile in a given subject as a weighted sum of local response profiles in the common space.

The searchlight hyperalignment algorithm generates an abstract feature space that does not directly map onto the anatomical space of any particular subject. This means we cannot directly compare, vertex by vertex, data in the common space generated by hyperalignment to data in any individual subject’s anatomical space. To directly compare the hyperaligned between-subject model to the other two types of models, for each leave-one-subject-out cross-validation fold we first transformed the training data—three runs for each of 17 subjects—into the common space using each subject’s unique hyperalignment transformation, and then mapped all 17 training subjects’ response vectors from the common space into the left-out test subject’s anatomical space (normalized to the fsaverage6 template) using the transpose of the left-out test subject’s hyperalignment transformation matrix. That is, for each left-out test subject, we mapped responses for the 17 training subjects into the left-out subject’s space *via* the common space. We then averaged response time series across training subjects. We did not apply any hyperalignment transformations to the validation data (the left-out test subject’s left-out test run). Note, however, that the whole-cortex matrix of local transformations learned by the searchlight hyperalignment algorithm is not orthogonal. This approach allows us to directly compare the three types of vertex-wise models on a subject-by-subject basis (i.e., when performing paired statistical tests). Whole-brain hyperalignment and several of the subsequent analyses were implemented using PyMVPA (Hanke et al., 2009).

### Semantic features

The *Life* documentary was annotated with a list of words describing the agents (i.e., animals), actions, objects, and scene for each camera angle of the movie. For example, if one camera angle depicted a giraffe eating grass on the savannah, the corresponding annotation would be the list of words “giraffe”, “eating”, “grass”, and “savannah”. Then, the camera angle annotations were interpolated for every 2.5 s of the movie, so that every imaging volume was assigned semantic labels. The annotation contained 277 unique words in total, and each imaging volume was assigned on average 5.28 words (*SD* = 1.91).

Next, we assigned a 300-dimensional word2vec semantic feature vector to each label in the annotation. We used pre-trained word2vec embeddings comprising a vocabulary of 3 million words and phrases trained on roughly 100 billion words from the Google News text corpus using the skip-gram architecture (Mikolov et al., 2013). Semantic vectors for all labels assigned to a given imaging volume were averaged to create a single 300-dimensional semantic vector per volume (cf. Vodrahalli et al., 2017). Mikolov and colleagues (2013) demonstrated that the word representations learned by the skip-gram model exhibit a linear structure that makes it possible to meaningfully combine words by an element-wise addition of their vector representations.

To accommodate the delayed hemodynamic response, we concatenated semantic vectors from the previous TRs (2.5, 5.0, 7.5, and 10.0 s; similarly to, e.g., Huth et al., 2012). The final vector assigned to each imaging volume for training and testing the encoding model comprised a concatenated 1,200-dimensional vector capturing the semantic content of the four preceding time points.

### Regularized regression

We estimated vertex-wise forward encoding models using the L2-penalized linear least squares regression (i.e., ridge regression) in three different ways: (*a*) within-subject models; (*b*) between-subject models using anatomical normalization; and (*c*) between-subject models using hyperalignment following anatomical normalization. All models were evaluated using leave-one-run-out cross-validation. In each of these leave-one-out runs, the within-subject and between-subject models were trained as follows: within-subject models were trained on three of the four imaging runs, then tested on the left-out fourth run separately for each subject. Between-subject models were trained on the averaged time series of 17 of the 18 participants over three of the four runs. The estimated between-subject models were then tested on the left-out fourth run in the left-out 18th participant. This yielded 72 total data folds for each of the three types of models. Figure 1 schematically depicts our approach for constructing between-subject semantic encoding models using hyperalignment.

**Figure 1.**
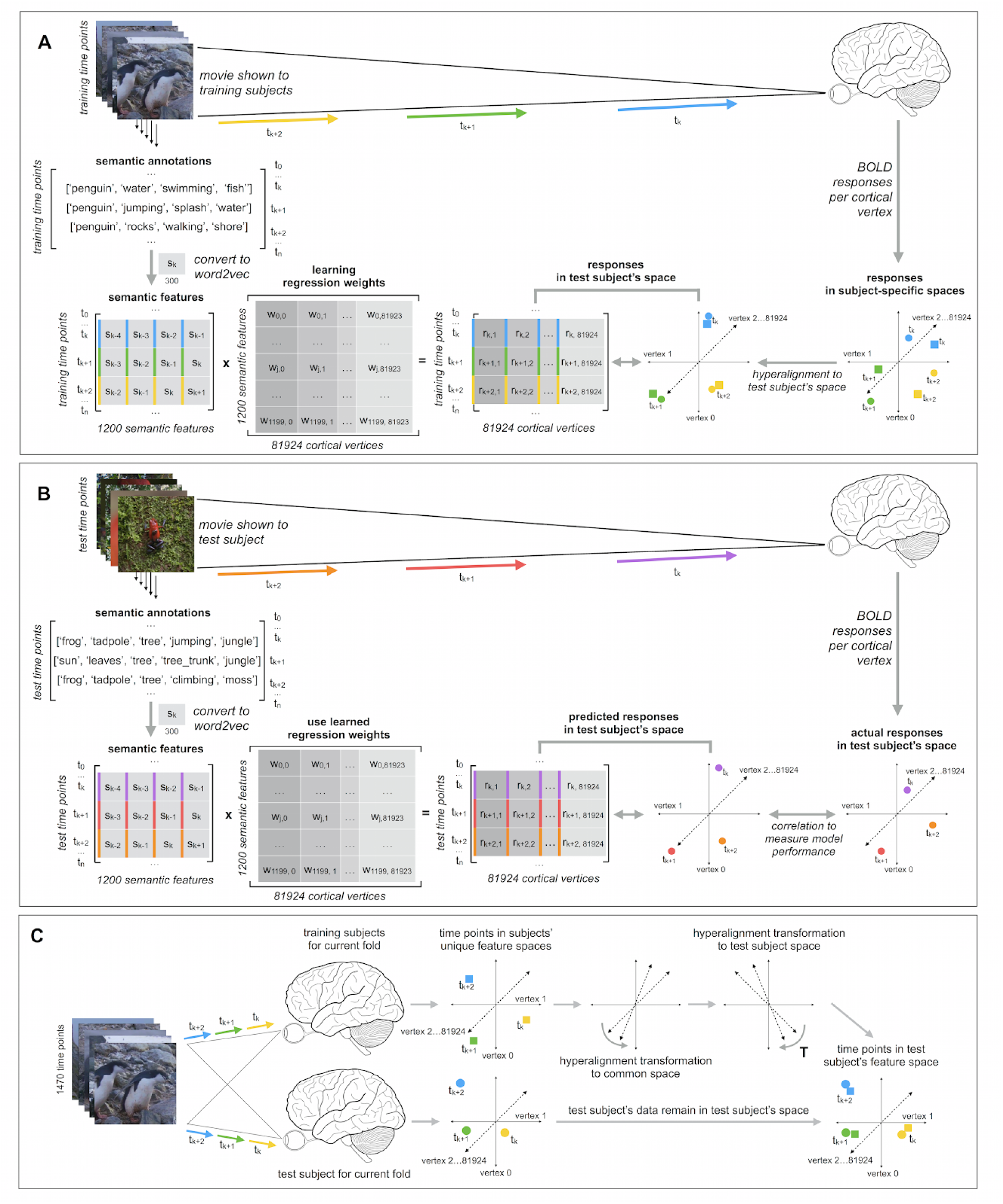
Schematic for constructing between-subject semantic encoding models using hyperalignment. The schematic depicts one fold of the nested leave-one-out cross-validation procedure repeated for four test runs and 18 test participants (72 cross-validation folds in total). (**A**) Training between-subject semantic encoding models using ridge regression. Regression coefficients (weights) are estimated to predict response time series per vertex based on three training runs. (**B**) Testing semantic encoding models. Regression weights estimated on training data to are used to predict response time series for a fourth test run. Model prediction performance is evaluated by computing the Pearson correlation between the predicted responses andare the actual response time series per vertex. (**C**) Hyperalignment for between-subject semantic encoding models. For each test subject in the leave-one-subject-out cross-validation procedure, we first projected each training subject’s data into the common space using their subject-specific hyperalignment transformations. We then use the transpose of the test subject’s hyperalignment transformation to project all training subjects’ data into the test subject’s space. We averaged response vectors for all training subjects in the test subject’s space, then trained the encoding model on this averaged response trajectory. Finally, we evaluated between-subject model performance by predicting vertex-wise response time series for the left-out test run in the left-out test participant, and computed the Pearson correlation between the predicted time series and the actual time series per vertex.

In these models, the number of predictor variables (1,200) exceeds the number of observations (ranging from 1,097 to 1,163 imaging volumes). We used ridge regression to estimate regression coefficients (weights) for the semantic predictor variables so as to best predict the response time series at each vertex. We used a modified implementation of ridge regression authored by Huth and colleagues (2012). Ridge regression uses a regularization hyperparameter to control the magnitude of the regression coefficients, where a larger regularization parameter yields greater shrinkage and reduces the effect of collinearity among predictor variables. The regularization parameter was chosen using leave­one­run­out cross­validation nested within each set of training runs. We estimated regression coefficients for a grid of 20 regularization parameters log­spaced from 1–1,000 at each vertex within each set of two runs in the training set of three runs. We then predicted the responses for the held­out third run (within the training set) and evaluated model prediction performance by computing the correlation between the predicted and actual responses. These correlations were averaged over the three cross­validation folds nested within the training set, then averaged across all vertices. We then selected the regularization parameter with the maximal model performance across runs and vertices. Selecting a single regularization parameter across all vertices ensures that estimated regression coefficients are comparable across vertices. This regularization parameter was then used at the final stage when estimating the encoding model across all three training runs for evaluation on the left­out fourth run. Note, however, that different regularization parameters were chosen for each of the four leave­one­run­out cross­validation folds (where both stimuli and hyperalignment transformation differed for each set of training runs), and for each of the 18 leave­one­subject­out cross­validation folds used for between­subject models. For the two between­subject models, the optimal regularization parameter was either 12.74 or 18.33 for every test subject (due to averaging response time series across training subjects). Note that these regularization parameters are considerably lower than those reported by Huth and colleagues (2016). This may be due to several factors, including our procedure for averaging time series across subjects during training, having fewer time points in the training set, and our use of the relatively dense lower­dimensional word2vec embeddings. However, for within­subject models, the optimal regularization parameter were more variable, likely due to increased noise.

To evaluate the vertex-wise forward encoding models, we used the regression coefficients from the model trained on three training runs to predict the response time series for the left-out fourth run. For between-subject models, we used the regression coefficients estimated on the training runs in the training subjects (transformed into the test subject’s space via the common space estimated using hyperalignment) to predict responses for the left-out run in the left-out subject. For between-subject models, both the hyperalignment transformations and encoding models were cross-validated to previously unseen data; the test run in the test subject played no role in estimating the hyperalignment transformations or the regression weights of the encoding model. For each vertex, we then computed the Pearson correlation between the predicted time series and actual time series for that run to measure model prediction performance (as in, e.g., Huth et al., 2012). Pearson correlations were Fisher *z*-transformed prior to statistical tests. We then averaged together the Pearson correlations for each of the four held-out test runs for visualization.

## Results

### Inter-subject correlations

To ensure that the common space learned by hyperalignment finds common bases for fine-grained functional topographies across subjects, we computed ISCs for both vertex-wise response time series and searchlight representational geometries using anatomical normalization and hyperalignment. To assess how hyperalignment impacts time series ISCs (Hasson et al., 2004), for each run of the movie we computed the correlations between each subject’s time series per surface vertex and the average of all other subjects before and after hyperalignment (Guntupalli et al., 2016). We then averaged these ISCs across all four movie runs; this results in a correlation for each subject for each vertex. We visualized the mean correlation across subjects for each vertex. Hyperalignment improved inter-subject correlations of time series. Figure 2A shows cortical maps of time series ISCs before and after hyperalignment. Hyperalignment increased the mean ISC of time series across vertices from 0.077 to 0.151.

**Figure 2.**
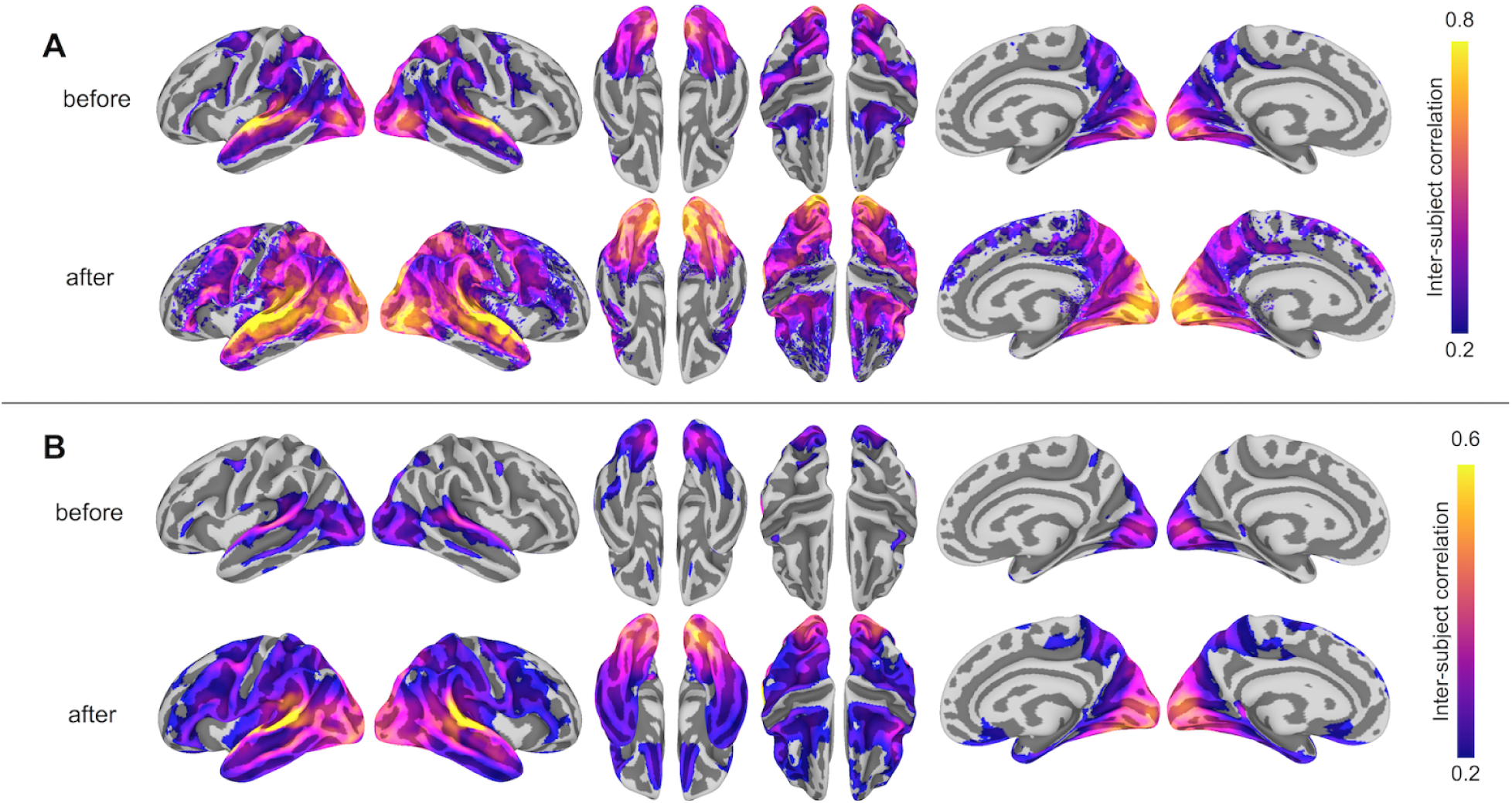
Hyperalignment improves inter-subject correlation (ISC) of response profiles and representational geometry. (**A**) ISC of vertex-wise response time series before and after hyperalignment. Colored vertices reflect the mean ISC across subjects, thresholded at a mean correlation of 0.2. ISCs are highest in the superior temporal gyrus (in the vicinity of auditory cortex), as well as the dorsal and ventral visual pathways, comprising early visual, lateral occipitotemporal, ventral temporal, posterior parietal and intraparietal cortices. (**B**) ISC of searchlight representational geometries (time-point representational dissimilarity matrices in 9 mm radius searchlights) before and after hyperalignment. Colored vertices reflect the mean pairwise correlation across subjects for each searchlight, thresholded at a mean correlation of 0.2, revealing a broader extent of cortex with improved alignment of functional topography after hyperalignment. ISCs were Fisher *z*-transformed before averaging across all subjects and inverse Fisher transformed before mapping onto the cortical surface for visualization. All maps are rendered on the fsaverage6 surface template.

We next analyzed the ISC of local representational geometries by calculating representational dissimilarity matrices (RDMs) comprising pairwise correlations between response vectors for all time points (Kriegeskorte et al., 2006; Oosterhof et al., 2011) in the test run of the movie using 9 mm radius surface-based searchlight disks (Kriegeskorte et al., 2006; Oosterhof et al., 2011). This procedure was repeated for each of the four runs. We averaged all pairwise correlations in the upper triangle of this matrix as well as averaging across runs. All operations involving correlations were performed after Fisher *z*-transformation and the results were inverse Fisher transformed for visualization. Hyperalignment also improved inter-subject correlations of searchlight representational geometries. Figure 2B shows cortical maps of ISCs of representational geometries before and after hyperalignment. Hyperalignment increased mean ISC of representational geometries across vertices from 0.157 to 0.230.

### Differences in model performance

We formally compared three types of semantic encoding models: within-subject models, between-subject models using anatomical normalization, and between-subject models using hyperalignment. For each subject, the within-subject model was compared to the between-subject models where that subject served as the test subject. For the hyperaligned between-subject model, group data were projected into the test subject’s space prior to model estimation. Figure 3 depicts model prediction performance for the three model types in two representative subjects, while Figure 4 depicts average model performance across subjects.

**Figure 3.**
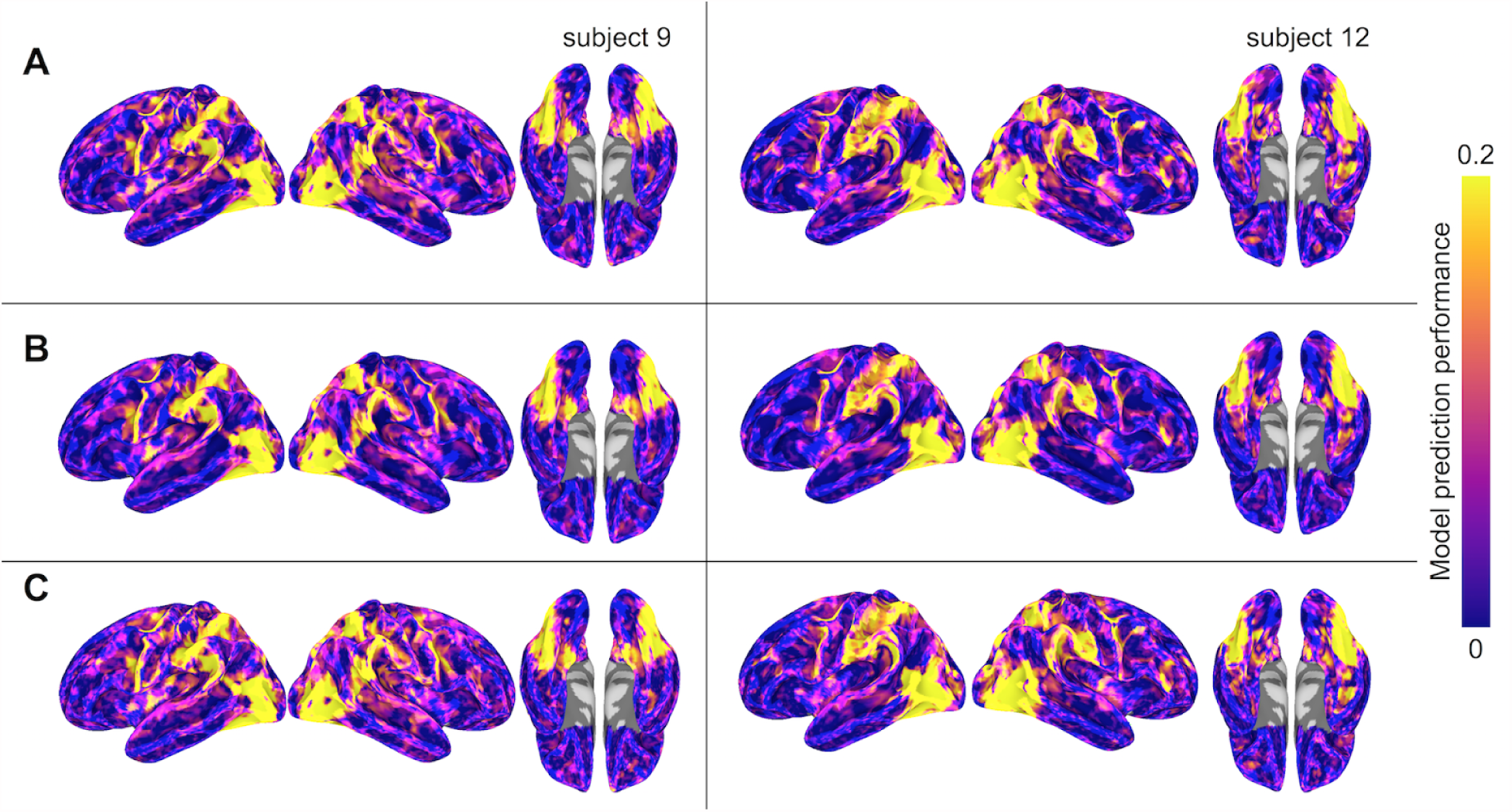
Model prediction performance maps for two representative subjects (left and right). Colored vertices reflect the Pearson correlation between the predicted and actual time series averaged across the four test runs. Three types of models are presented: (**A**) within-subject model performance maps; (**B**) between-subject anatomically aligned model performance maps; and (**C**) between-subject hyperaligned model performance maps. All maps are unthresholded and uncorrected for multiple tests.

**Figure 4.**
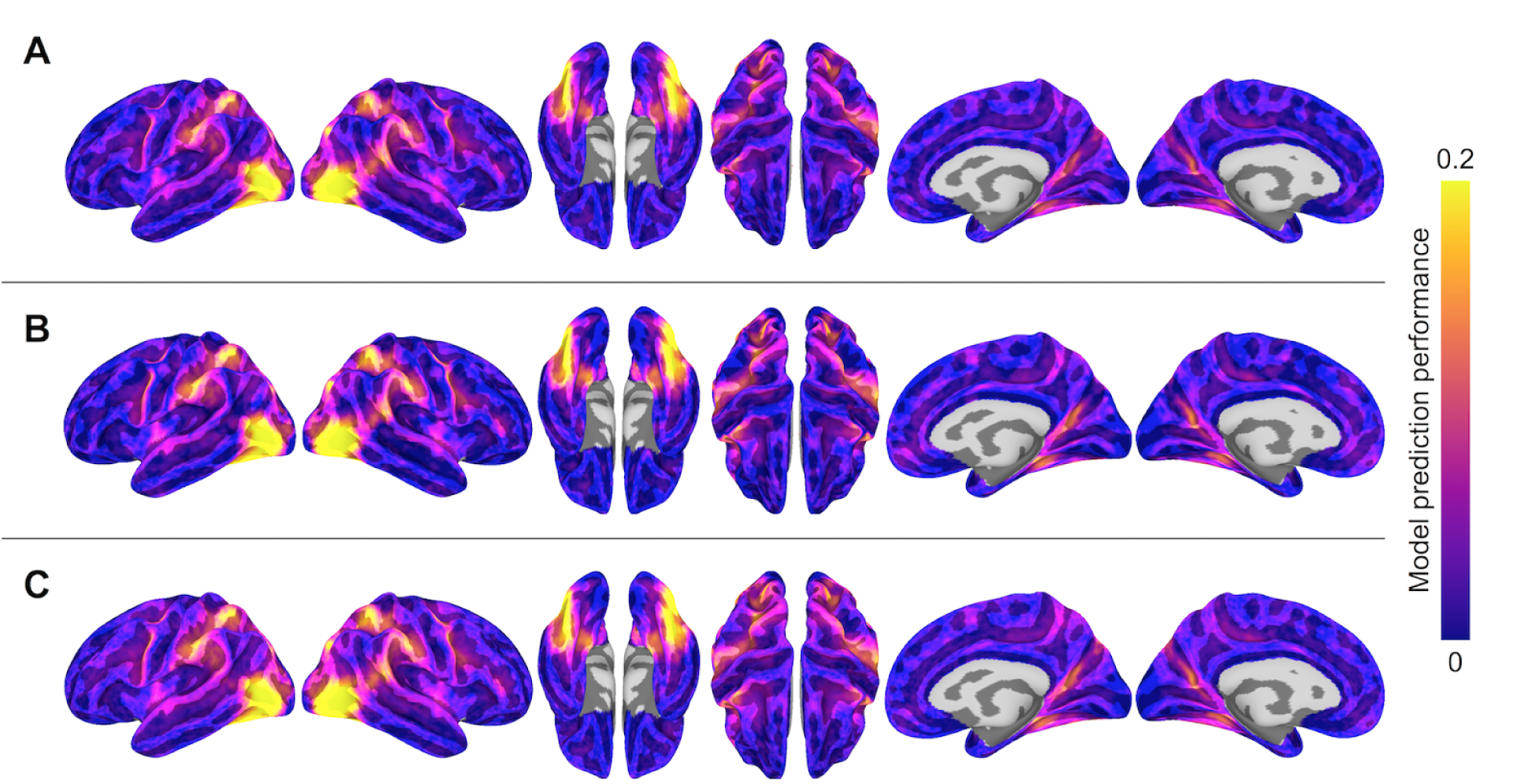
Model prediction performance maps averaged across subjects. Colored vertices reflect the Pearson correlation between the predicted and actual time series averaged across the four test runs and averaged across subjects. Three types of models are presented: (**A**) within-subject model performance maps; (**B**) between-subject anatomically aligned model performance maps; and (**C**) between-subject hyperaligned model performance maps. All maps are unthresholded and uncorrected for multiple tests.

We summarized differences in model performance across the entire cortex in two ways. To constrain our analysis to well-predicted vertices, for each subject we selected the 10,000 vertices with highest model performance separately for each model. We then considered only the union of well-predicted vertices across all three models (on average 15,436 vertices per subject, *SD* = 1,196 across subjects). First, for each pair of models, we computed the proportion of vertices with greater model prediction performance (i.e., correlation between predicted and actual time series for the test data) for one model relative to the other. We calculated these proportions per subject, then computed a paired *t*-test to assess statistical significance per model pair. When comparing the model performance for the within-subject and the between-subject models, the between-subject model using anatomical alignment yielded higher correlations in 50.3% of selected cortical vertices (*t*(17) = 0.307, *p* = 0.762). The between-subject model using hyperalignment yielded better performance than the within-subject model in 58.8% of selected cortical vertices (*t*(17) = 8.486, *p* < 0.001). The between-subject model using hyperalignment also yielded better performance than the between-subject model using anatomical alignment (59.3% of cortical vertices; *t*(17) = 20.200, *p* < 0.001).

Second, we assessed the difference in model prediction performance averaged across the same subset of well-predicted vertices. The between-subject model using anatomical alignment performed similarly to the within-subject model (0.127 and 0.131, respectively; *t*(17) = 1.851, *p* = 0.082). The between-subject model using hyperalignment performed better than the within-subject model (0.143 and 0.131, respectively; *t*(17) = 7.384, *p* < 0.001). Additionally, hyperalignment exceeded anatomical alignment when comparing the performance of between-subject models (0.143 and 0.127, respectively; *t*(17) = 16.185, *p* < 0.001).

To visualize differences in model performance, we compared model performance maps on the cortical surface (Figure 5). We computed vertex-wise paired *t*-tests for each of the three model comparisons. For visualization, we thresholded maps at a *t*-value of 2.11 (*p* < .05, two-tailed test, uncorrected for multiple tests).

**Figure 5.**
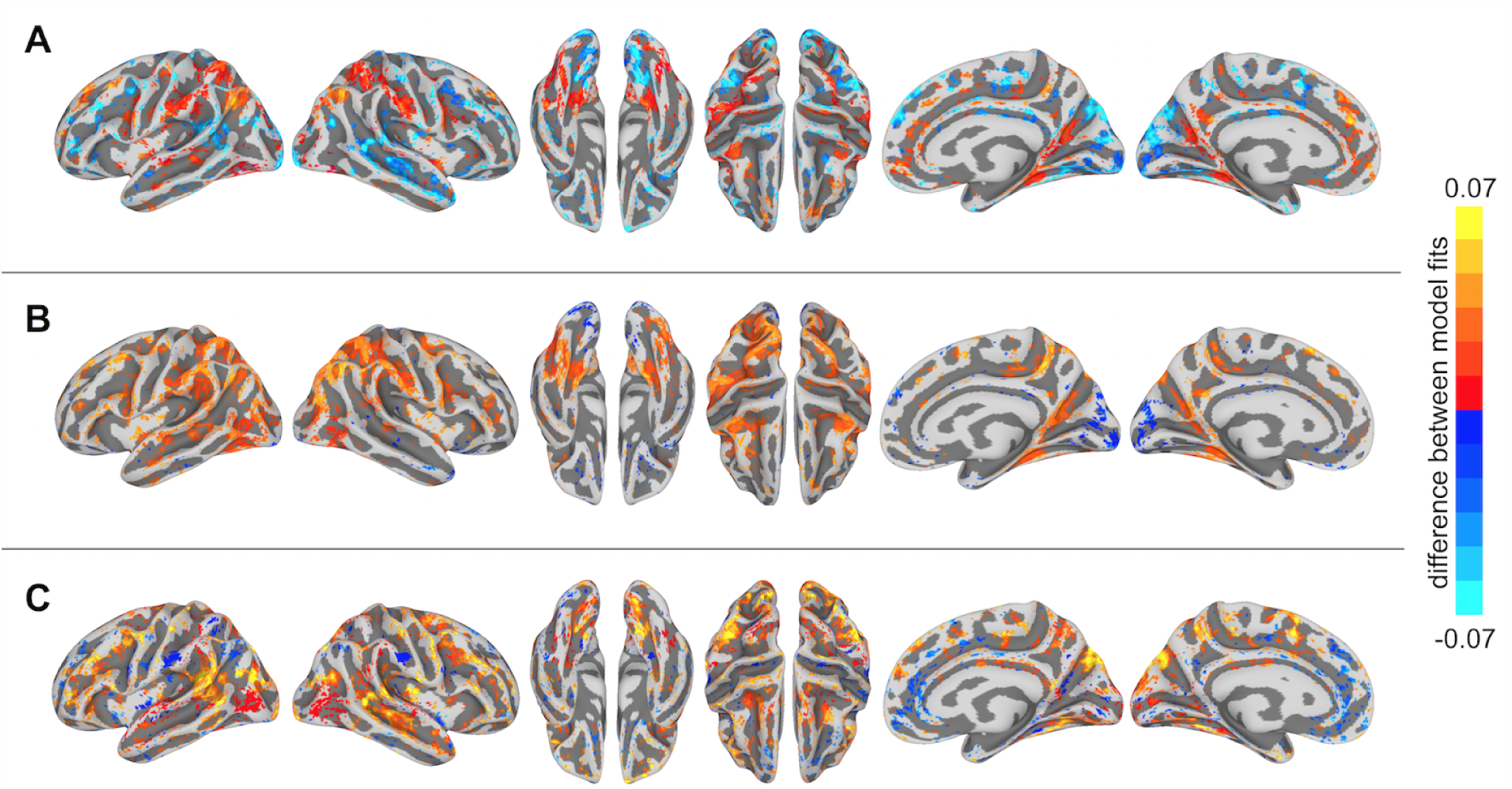
Differences in semantic encoding model performance maps. (**A**) Paired differences in model performance between between-subject models using anatomical normalization and within-subject models. Warm colors indicate vertices where the anatomically-aligned between-subject model performance exceeds within-subject model performance, and cool colors indicate where within-subject model performance exceeds anatomically-aligned between-subject model performance. (**B**) Paired differences in model performance between between-subject models using hyperalignment and within-subject models. Warm colors indicate vertices where the hyperaligned between-subject model performance exceeds within-subject model performance, and cool colors indicate where within-subject model performance exceeds hyperaligned between-subject model performance. (**C**) Paired differences in model performance for between-subject models using hyperalignment and anatomical normalization. Warm colors indicate vertices where the hyperaligned between-subject model performance exceeds anatomically-aligned between-subject model performance, and cool colors indicate where anatomically-aligned between-subject model performance exceeds hyperaligned between-subject model performance. Colored vertices reflect mean paired differences in model performance, thresholded at an absolute *t*-value of *t*(17) = 2.11, *p* < .05, uncorrected for multiple comparisons.

### Spatial specificity of semantic tuning

To compare the spatial specificity of semantic tuning across model types, we computed the spatial point spread function (PSF) of the semantic model predictions (Figure 6). To constrain our analysis to well-predicted vertices, for each subject we again selected the 10,000 vertices with highest model performance separately for each model and considered only the union of these well-predicted vertices across all three models. For each well-predicted vertex, we computed the model prediction performance (Pearson correlation between predicted and actual time series) for that vertex, and for neighboring vertices using the same prediction equation at 2 mm intervals up to 12 mm. That is, we used the encoding model at each vertex to predict the actual time series at neighboring, increasingly distant vertices. Each “ring” of vertices (e.g., the ring of vertices at a radius 10–12 mm from the central vertex of interest) was 2 mm wide and excluded vertices sampled at smaller radii. For a given ring of vertices, model performance was computed at each vertex in the ring and averaged across those vertices. Model performances at each radius per vertex were then averaged across the set of selected well-predicted vertices. To statistically assess PSFs, we computed bootstrapped confidence intervals around the model performance estimates at each radius by resampling subjects with replacement. To quantify the decline in spatial specificity of model performance over radii, we fit a logarithmic function to the PSF for each model at the midpoint of each ring (i.e., the vertex of interest, 1 mm, 3 mm, etc.) and reported the slope of this fit. The spatial point-spread function of the model predictions for the between-subject model using anatomical alignment was relatively flat (negative slope of the logarithmic fit = 0.0230 [0.0222, 0.0237]), reflecting spatially smooth semantic tuning. By contrast, the within-subject and hyperaligned between-subject models have steeper slopes (0.0424 [0.0397, 0.0453] and 0.0420 [0.0404, 0.0437], respectively; both p < 0.001), indicating greater spatial specificity in semantic tuning.

**Figure 6.**
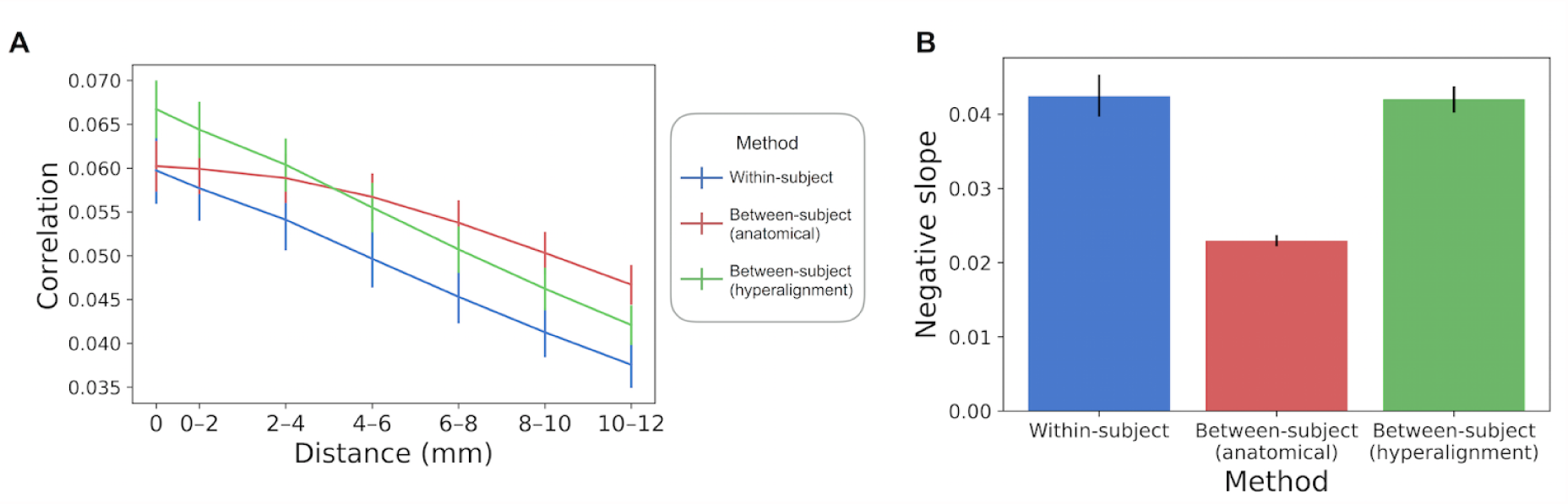
Spatial point spread function of semantic tuning. (**A**) Correlation between the predicted time series of one vertex and the actual time-series of its neighboring vertices up to 12 mm away. Correlations were aggregated based on distance from the central vertex of interest and averaged across vertices and subjects. Error bars denote 68% confidence intervals (standard error of the mean). (**B**) The within-subjects and hyperaligned between-subject models have the steepest slopes (negative of the slope based on logarithmic curve fitting). Error bars denote 95% confidence intervals obtained by bootstrapping subjects 20,000 times with replacement.

The prediction performance maps for each model varied in their spatial smoothness (Figure 7). We computed the full width at half maximum (FWHM) of the model performance maps using SUMA’s SurfFWHM. Spatial smoothness was computed per run in each hemisphere in each participant and averaged across hemispheres. Model performance maps for the between-subject model using anatomical alignment were significantly more spatially blurred than for the within-subject model (5.043 mm FWHM and 4.439 mm FWHM, respectively; *t*(17) = 26.807, *p* < 0.001). The between-subject model using hyperalignment recovered the spatial specificity of the within-subject maps, and in fact yielded less smooth model performance maps (3.711 mm FWHM) than the within-subject model (*t*(17) = 21.451, *p* < 0.001).

**Figure 7.**
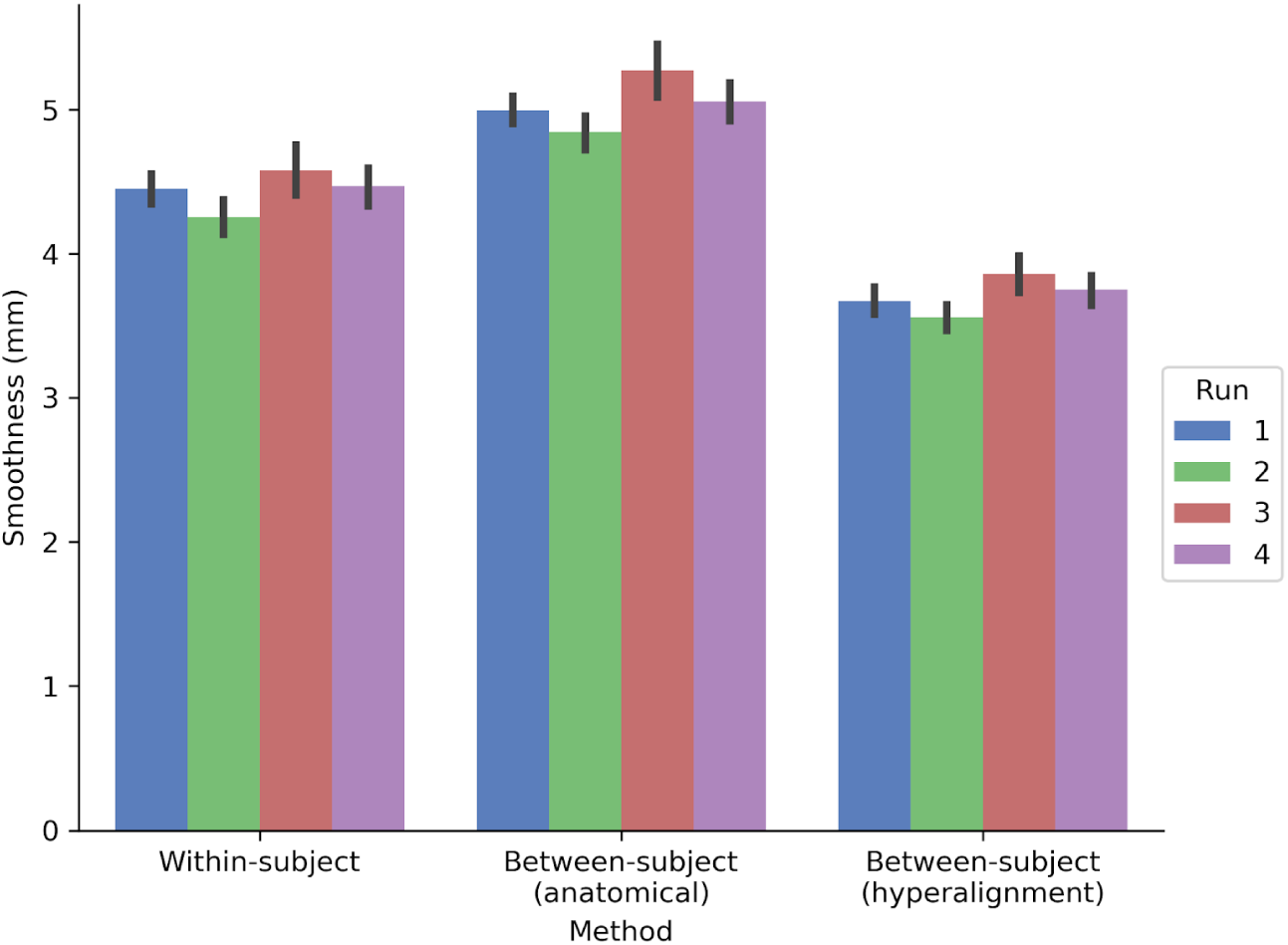
Spatial smoothness (FWHM) of model prediction performance maps on the cortical surface. The between-subject model performance maps using anatomical alignment are blurred relative to the within-subject model performance maps, while the hyperaligned between-subject model recovers the spatial specificity of the within-subject model. The height of each bar indicates spatial smoothness averaged across hemispheres and participants for each run. Error bars indicate bootstrapped 95% confidence intervals estimated by resampling participants (1,000 bootstrap samples).

We also assessed how well the between-subject model performance maps approximated the spatial organization of the within-subject model performance maps by computing the Pearson correlation between model performance maps (Figure 8). Correlations were computed across both cortical hemispheres within each participant and run. The spatial correlation between the model performance maps for the within-subject and between-subject models was .544 using anatomical normalization and .721 when using hyperalignment. That is, the spatial correlation between the map of within-subject model fits and the map of between-subject model fits increased by .177 after hyperalignment (a 33% increase; *t*(17) = 22.454, *p* < 0.001).

**Figure 8.**
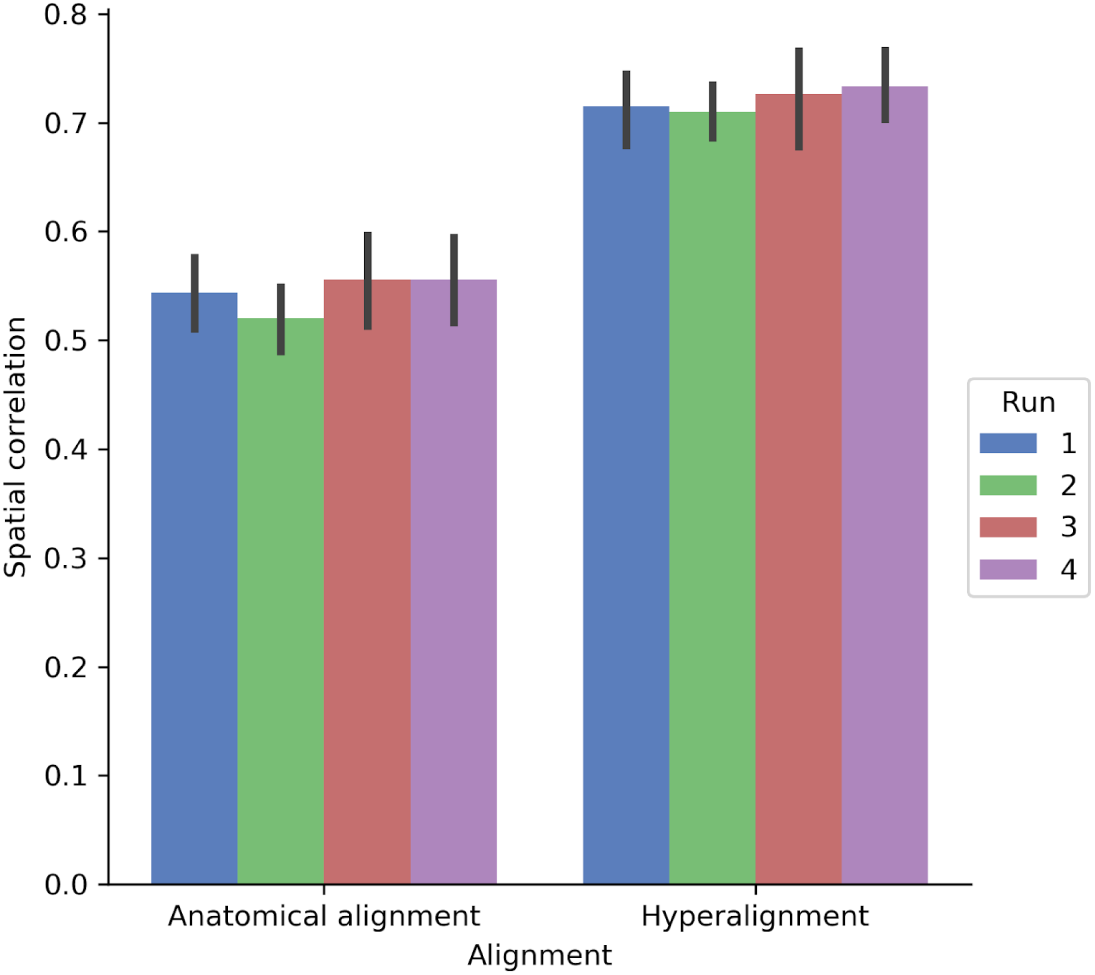
Spatial correlation between within-subject model prediction performance maps and between-subject model prediction performance maps using either anatomical alignment or hyperalignment. The between-subject model using hyperalignment yielded a model performance map that is more similar to the within-subject model than the model performance map of the between-subject model using anatomical alignment. The height of each bar indicates mean spatial correlation across participants for each run. Error bars indicate bootstrapped 95% confidence intervals estimated by resampling participants (1,000 bootstrap samples).

## Discussion

We developed a framework for constructing between-subject semantic encoding models that generalize to both novel stimuli and novel subjects. Vertex-wise forward encoding models were used in conjunction with hyperalignment to translate fine-grained functional topographies across individuals. Naturalistic experimental paradigms that broadly sample neural representational space play a critical role in this procedure, effectively enhancing the generalizability of both the encoding model and the hyperalignment transformations (Haxby et al., 2011, 2014).

Typically, encoding models are estimated separately for each subject using a relatively large volume of data (e.g., Huth et al., 2012; Mitchell et al., 2008; Pereira et al., 2018). Mirroring recent reports on resting-state functional connectivity in highly-sampled individuals (Gordon et al., 2017; Laumann et al., 2015), these within-subject models can reveal highly detailed, idiosyncratic functional organization. However, there is a trade-off: we can only acquire large volumes of data in relatively few individuals (often the authors themselves, e.g., Gordon et al., 2017; Huth et al., 2012, 2016; Laumann et al., 2015; Nishimoto et al., 2011). This inherently limits the generality of conclusions drawn from within-subject models and undercuts efforts to relate the acquired data to between-subject variables. Constructing between-subject models that make individualized predictions in novel subjects is a critical step toward increasing the utility of cognitive neuroscience (Gabrieli et al., 2015). Although between-subject models can be constructed using anatomical normalization, this obscures considerable heterogeneity in functional organization because fine-scale variations in functional tuning are not tightly tethered to macroanatomical features (Guntupalli et al., 2016, 2018). Hyperalignment affords aggregation of data across individuals that aligns these fine-scale variations, thus alleviating this tension. In constructing a common representational space, we decouple functional tuning from anatomy, registering representational geometries rather than anatomical features. Unlike anatomical normalization, averaging across subjects in this space does not collapse responses that map onto topographies that are idiosyncratic to individual brains. Critically, we also preserve each individual’s idiosyncratic functional–anatomical mapping in their respective transformation matrix, allowing us to project group data into any individual subject’s anatomical space with high fidelity. This precision mapping enables out-of-sample prediction on the scale of individual voxels (Dubois and Adolphs, 2016; Poldrack, 2017).

Overall, our findings demonstrate that between-subject models estimated using hyperalignment outperform within-subject models. Between-subject models estimated using anatomical normalization yield artificially smooth maps of semantic tuning. Hyperalignment, on the other hand, retained the spatial specificity of within-subject models. The semantic encoding model used here best predicted responses in a network of areas previously implicated in representing animal taxonomy (Connolly et al., 2012, 2016; Sha et al., 2015) and observed action (Nastase et al., 2017; Oosterhof et al., 2013; Wurm et al., 2016; Wurm and Lingnau, 2015), including ventral temporal, lateral occipitotemporal, anterior intraparietal, and premotor cortices. Interestingly, inter-subject correlations were highest in superior temporal cortex, encompassing auditory areas. Although this suggests that the auditory narrative evoked highly reliable neural responses, in the present analyses the linguistic content of the narrative was not explicitly included in the semantic annotation.

The current approach for constructing between-subject encoding models using hyperalignment differs in several ways from related reports. Yamada and colleagues (2015) introduced a sparse regression algorithm for predicting voxel responses across pairs of subjects. This algorithm estimates a more flexible mapping than the Procrustes transformation between pairs of subjects and does not yield a common representational space across all subjects. Their approach was evaluated in early visual cortex using 10 pixel × 10 pixel black-and-white geometric images. While more recent work by Güçlü and van Gerven (2017) used naturalistic visual stimuli, subjects in this study were required to perform a highly non-naturalistic central fixation task (Nishimoto et al., 2011). Both Yamada et al. (2015) and Güçlü and van Gerven (2017) validated their models in a limited cohort of three subjects. Vodrahalli and colleagues (2017) used a variant of hyperalignment to estimate encoding models in a lower-dimensional (20 dimensions) common space. Models were evaluated in this low-dimensional shared space using a scene classification analysis. Their findings suggest that using a weighted averaging scheme for aggregating word embeddings assigned to a given imaging volume can improve model performance. However, in that experiment, annotators provided natural-language descriptions of the film (including many uninformative words). In the current study, the annotation included only the most salient or descriptive labels, effectively filtering out stop words and otherwise uninformative labels. Unlike previous reports, which limited their analyses to one or several regions of interest, we used searchlight hyperalignment to derive a locally-constrained common space for each cortical hemisphere and estimated between-subject encoding models across the entire cortex. Finally, none of the previously mentioned studies projected group data (via the common space) into each test subject’s idiosyncratic response space prior to model estimation. Here, we used data from a naturalistic stimulus and task, transformed through a high-dimensional common space into each subject’s idiosyncratic anatomical space, to estimate between-subject models across all cortical vertices.

Although the current findings demonstrate the utility of hyperalignment in constructing between-subject encoding models, there are several open questions. Under what circumstances will a between-subject model outperform within-subject models? Averaging group data in a common representational space provides a more robust estimate of response trajectories without sacrificing anatomical specificity. This should provide an advantage when predicting semantic tuning for noisy features in a given test subject, as the group estimate will be more robust. In addition to leveraging a larger volume of group data with the precision of within-subject models, hyperalignment effectively filters response profiles, suppressing variance not shared across subjects (Guntupalli et al., 2018). More generally, between-subject models can improve performance in areas where responses are highly stereotyped across individuals. For example, in the current study, both types of between-subject models improved model performance in anterior intraparietal areas, which are implicated in observed action representation during natural vision (Nastase et al., 2017).

When will hyperalignment fall short of within-subject performance? First, within-subject performance should be superior by virtue of capturing idiosyncratic functional tuning, but it is usually impractical or impossible to collect sufficient data in each subject. Because hyperalignment largely preserves each subject’s representational geometry, we expect any advantage will be attenuated when the test subject’s representational geometry is idiosyncratic, irrespective of functional–anatomical correspondence (Charest et al., 2014; Kriegeskorte and Kievit, 2013). Note, however, that hyperalignment may serve to disentangle idiosyncrasies in representation from idiosyncrasies in functional–anatomical correspondence. Furthermore, we would not expect an advantage from hyperalignment if the stimulus or experimental paradigm used to derive the hyperalignment transformations did not adequately sample the neural representational subspaces important for estimating the encoding model. This concern becomes relevant if, in contrast to the current study, hyperalignment parameters and the encoding model are estimated on data derived from experimental paradigms that are more restricted and non-naturalistic. Finally, representations that are encoded in a coarse-grained or anatomically stereotyped manner will benefit less from hyperalignment, and anatomical normalization may be sufficient. However, as the resolution and sensitivity of functional measurements improves, and as more sophisticated encoding models begin to make finer-grained predictions, hyperalignment will become increasingly necessary.

## Author Contributions

CEVU, SAN, ACC, MF, MIG, and JVH designed the experiment; SAN and ACC collected the data; CEVU, SAN, MF, and IH analyzed the data; CEVU, SAN, ACC, MF, and JVH wrote the manuscript.

## Acknowledgments

We thank Matteo Visconti di Oleggio Castello, Yaroslav O. Halchenko, Vassiki Chauhan, Easha Narayan, Kelsey G. Wheeler, Courtney Rogers, and Terry Sackett for helpful suggestions and administrative support.

